# Abemaciclib restricts HCMV replication by suppressing pUL97-mediated phosphorylation of SAMHD1

**DOI:** 10.1101/2023.02.01.526617

**Authors:** Georgios Vavouras Syrigos, Maximilian Feige, Alicia Dirlam, Ramona Businger, Iris Gruska, Lüder Wiebusch, Klaus Hamprecht, Michael Schindler

## Abstract

Human cytomegalovirus (HCMV) is a herpesvirus that causes life-threatening infections in newborns or immunosuppressed patients. For viral replication, HCMV establishes a network of cellular interactions, among others cyclin-dependent kinases (CDK). Furthermore, HCMV encodes pUL97, a viral kinase, which is a CDK-homologue. HCMV uses pUL97 in order to phosphorylate and thereby antagonize SAMHD1, an antiviral host cell factor. Since HCMV has several mechanisms to evade restriction by SAMHD1, we first analyzed the kinetics of SAMHD1-inactivation and found that phosphorylation of SAMHD1 by pUL97 occurs directly after infection of macrophages. We hence hypothesized that inhibition of this process qualifies as efficient antiviral target and FDA approved CDK-inhibitors (CDKIs) might be potent antivirals that prevent the inactivation of SAMHD1. Indeed, Abemaciclib, a 2^nd^ generation CDKI exhibited superior IC50s against HCMV in infected macrophages and the antiviral activity largely relied on its ability to block pUL97-mediated SAMHD1-phosphorylation. Altogether, our study highlights the therapeutic potential of clinically-approved CDKIs as antivirals against HCMV, sheds light on their mode of action and establishes SAMHD1 as a valid and highly potent therapeutic target.

## INTRODUCTION

Human cytomegalovirus (HCMV) is the major congenital infection with severe consequences for the neonates, and is transmitted from the mother to the fetus during pregnancy^1^. HCMV infection can cause a variety of symptoms in newborns, like mental and growth retardation, motor difficulties, microcephaly, hearing loss and other^2,3^. In addition, HCMV is a threat for immunocompromised individuals, like AIDS patients and organ transplant recipients, and a primary infection or a viral re-activation can have devastating effects, like increased morbidity and mortality^4,5^. HCMV has a large genome of 235 kb, which encodes for more than 700 translated open reading frames and microRNAs^6–8^. It has been co-evolved with humans for thousands of years, therefore it has developed multiple sophisticated mechanisms to escape the human immune system^9^ and develop lifelong latency. These characteristics of the virus make the development of antiviral compounds against HCMV challenging. There are some drugs available for prophylaxis or as a preemptive therapy^4,10^, but they have limitations as for instance nephrotoxicity, low bioavailability and emergence of drug-resistant strains^11^.

SAMHD1 is a host protein which among others possesses a dNTP-ase activity^12,13^, that is vital for cell division and modulated by cyclin dependent kinases (CDKs), specifically CDK1 and CDK2^14–20^. It has been shown that SAMHD1 bears antiviral activity against a panel of viruses^21^ (HIV-1^13,22–26^, HSV-1^27,28^, vaccinia virus^27^, HBV^16,20^, HCMV^29–31^) which might rely on its activity in depleting dNTPs. Nevertheless the hydrolase and antiviral activities have not been completely linked yet^14,16,19,22^. HCMV, in turn, antagonizes SAMHD1 using numerous different mechanisms including proteasomal degradation^29,31^, mRNA transcriptional suppression^29^, subcellular mislocalization^31^ and phosphorylation^16,29,30^. Phosphorylated SAMHD1 is antivirally inactive^14,16,20,29^, however it is still unclear, if pSAMHD1 retains its dNTP-ase activity^14,16,19,22^. Apart from the HCMV-encoded kinase pUL97, a viral CDK which has been shown to phosphorylate SAMHD1^16,29,30^, HCMV hijacks host cell CDKs by activating the CDK-cyclin complex^32–35^. This might contribute to effectively keep SAMHD1 in its inactive state.

Pharmacological CDKIs are compounds that impair the activity of CDKs, thus effecting the cell cycle, and they are developed and used as a treatment against various types of cancer^36–38^. Since viruses manipulate the cell cycle for efficient replication^34^, CDKIs have been studied as potential anti-viral treatment^17,20,39–45^. The FDA approved CDK4/6 inhibitor Palbociclib for instance, is on the one hand used against breast cancer, but on the other hand shown to inhibit HSV-1 and HIV-1^17,44,45^ infection in primary macrophages, while a CDK7 inhibitor, LDC4297 has antiviral activity against a broad range of herpesviruses, including HCMV^41,43^. Some combination of CDKIs with each other or with currently used anti-HCMV treatment exhibits synergistic effects^39,40^.

However, the mode of antiviral action of the CDKIs and the reason underlying their highly differential antiviral activity is yet unclear. We hypothesized, that the antiviral activity of CDKIs is related to their ability to suppress SAMHD1-phosphorlyation and thus convert it into its antiviral active form. We first unraveled the kinetics of SAMHD1 antagonism upon HCMV infection in primary macrophages and tested various CDKIs, including a series of clinically approved CDK4/6 inhibitors for their ability to inhibit HCMV replication and suppress SAMHD1-phosporylation. By this, we revealed the relevance of SAMHD1 as druggable target and the high potential of CDKIs for antiviral therapy.

## METHODS

### Cell culture

Primary human macrophages (MΦs) were cultured in macrophage medium (RPMI-1640 GlutaMAX^™^ (Thermo) supplemented with 4% human AB serum (Sigma), 100 μg/ml penicillin-streptomycin (Sigma), 1 mM sodium pyruvate (Gibco), 1x non-essential amino acids (Gibco) and 0.4x MEM vitamins (Sigma). Human foreskin fibroblasts (HFFs, ATCC SRC-1041) and ARPE-19 cell line (ATCC CRL-2302) were cultured in DMEM, high glucose, GlutaMAX™ (Thermo) supplemented with 5% fetal calf serum (FCS, Gibco) and 100 μg/ml penicillin-streptomycin. Human embryonic kidney (HEK) 293T cell line (DSMZ ACC635) was cultured in DMEM, high glucose, GlutaMAX™ (Thermo) supplemented with 10% FCS and 100 μg/ml penicillin-streptomycin. All cells were kept for growth in an environment of 37°C and 5% CO_2_.

### Isolation and differentiation of primary human macrophages

Monocyte derived macrophages (MDMs) were isolated from buffy coats of blood donors who gave informed consent for the use of blood-derived products for research purposes. We do not collect data concerning age and ethnicity, and we comply with all relevant ethical regulations approved by the ethics committee of the University Hospital Tübingen (IRB no. 507/2017B01). All buffy coat donations are received in pseudonymous form and chosen randomly. Peripheral blood mononuclear cells were separated from blood by Ficoll^®^ Paque Plus (Sigma) gradient centrifugation. Cells were differentiated by plastic adherence in macrophage medium for 4 days. Next, medium was changed to fresh MΦ medium, and cells were differentiated for 3 more days before using them for experimental procedures.

### Viruses and clinical isolates

The HCMV strain TB40/E was established after propagation of the clinical isolate TB40 in endothelial cells, which maintained the endothelial/macrophage tropism^46^. HCMV-delUL16-GFP was generated by a modification of the TB40/E after the replacement of UL16 with GFP^47^. The HCMV clinical isolates were isolated from urine and were kindly provided by Prof. Dr. Klaus Hamprecht and his group.

### HCMV stock and VLP generation

To generate fresh viral stocks, HFFs were infected with HCMV TB40/E for 7 to 10 days. At the same day of the planned infection of MDMs, the virus-containing media of infected HFFs was collected and centrifuged for 10 mins at 3,200g, to clear from cell debris. To produce VSV-G pseudotyped virus like particles (VLPs), either pSIV_Vpx- or pSIV_Vpx+ were co-transfected with pHIT60 (VSV-G) into HEK 293T cells, using a standard calcium phosphate method. Six hours post transfection, medium was changed to fresh HEK medium and supernatants containing the VLPs were collected 24 h later. VLP containing supernatants were centrifuged at 3,200g for 10 mins to clear from cell debris and instantly frozen at −80°C. VLP Vpx+ were tested for their efficiency to knock out SAMHD1 prior to use. 1.0E+05 MDMs per well were seeded in a 12-well plate and were treated with VLP Vpx+ or Vpx- or medium for 2 hours. After a medium change, cells were incubated for a total of 24 hours and then harvested, intra-stained for SAMHD1 and SAMHD1 levels were detected via flow cytometry.

### Cells infection

MDMs were seeded in a 6-well or 12-well plate (2.0E+05 or 1.0E+05 cells respectively) the day before infection and were infected with fresh virus stock. After 30 minutes of incubation, cells were spinoculated for 45 minutes, at 34°C, 600 g. After a subsequent incubation of 1 hour, media was changed to fresh MDM media and cells were incubated furthermore in cell culture conditions. HFFs were incubated with TB40/E or clinical isolates for 2 hours and after a medium change were furthermore incubated for a total of 24 hours.

### Flow cytometry staining

Cells were fixed with 2% paraformaldehyde (PFA) for 15 minutes at room temperature (RT), washed with PBS containing 1% FCS (FACS buffer) and permeabilized with 90% ice-cold methanol in water for 20 minutes at 4°C. Cells were then washed twice with FACS buffer and blocked with 10% FCS for 30 minutes at RT. Afterwards, the cells were stained with either rabbit SAMHD1 (ProteinTech, dilution 1:1000) or rabbit pSAMHD1 (specific for T592 phosphorylation, ProSci; dilution 1:1,000) antibody for 1 h at RT. After washing once, the cells were stained with goat anti-rabbit Alexa Fluor® 555 (Thermo, 1:1,000) and mouse anti-IE Alexa Fluor® 488 (Merck, 1: 1,000) for 1 hour at RT. Cells were washed as a final step and measured with the flow cytometer MACSQuant VYB (Miltenyi).

### Immunofluorescence staining

HFFs were washed once with PBS and fixed/permeabilized with 80% Acetone in water for 10 mins at RT. Subsequently cells were washed 3 times, blocked for 30 minutes at RT with 10% FCS in PBS, and stained with mouse anti-IE Alexa Fluor® 488 (Merck, 1:1,000) for 1 hour at RT. After a 3-times wash, nuclei were stained using DAPI in 1:20,000 dilution in PBS for 10 minutes at RT. A last washing step was conducted before taking the plates to the imaging multi-mode reader Cytation3 (Biotek).

### Compound treatment experiments

In the CDK4/6 inhibitor treatments, MDMs were pre-incubated with Palbociclib, Ribociclib or Abemaciclib (Selleckhem) for 16 hours before infection. After removing the virus, cells were furthermore inoculated with CDKis for 1,2,4 or 6 days. The positive control Ganciclovir (Selleckhem), as well as the panel of different CDKIs (K03861, JSH-150, LDC4297; Selleckhem) were added once, after the virus removal and incubated for 2 or 6 days.

### MTT assay

The MTT assay^48^ is a viability technique to determine the toxicity of the various compounds tested. In short, MDMs were seeded in 96-well plates and treated with the different compounds for 6 days. Cells were then washed and incubated in 10% MTT solution^48^ in phenol red-free medium for 3 hours at 37°C. Afterwards, medium was removed and the form crystals were dissolved in 0.04 M HCl in isopropanol in a plate shaker for 10 mins at RT. Absorption was measured at 570/650 nm using the Tristar^2^ S LB 942 Multimode Reader.

### In vitro kinase assay

In vitro kinase assays were performed as previously described^29^. Purified, His-tagged SAMHD1 served as substrate. HA-tagged pUL97 kinase was immuno-precipitated from transfected HEK 293T cells using the rat anti-HA antibody clone 3F10 (Roche). Kinase inhibitors were added to the kinase reactions at final concentrations of 1 μM (CDKIs) and 5 μM (Maribavir) respectively.

### qRT-PCR

RNA of cells was transcribed to cDNA using the QuantiTect Reverse Transcription Kit (Qiagen): 50-1000 ng RNA were mixed with RNAse-free water and gDNA Wipeout buffer for 2 min at 42°C. Afterwards reverse transcriptase-Master mix (Quantiscript Reverse Transcriptase + Quantiscript RT Buffer 5x + RT Primer Mix) was added and again incubated at 42°C for 15 min. Reverse transcriptase was deactivated by incubation at 95°C for 3 min and then samples were diluted in RNAse-free water. RT-qPCR was performed using the LightCycler®480 SYBR Green I Master-Ki (Roche): Primers were used for GAPDH (forward: TGCACCACCAACTGCTTAGC; reverse: GGCATGGACTGTGGTCATGAG) and SamHD1 (forward: CGAGATGTTCTCTGTGTTCA; reverse: CGTCCATCAAACATGTGAGA). Primers were mixed with PCR-grade water, before the addition of SYBR green Master mix. SYBR green-Master/Primer mix was added to cDNA-sample in qPCR-plates. qRT-PCR for each sample was performed in duplicates. After covering the plates and short centrifugation to combine the liquids qRT-PCR was then performed using the 480 LightCylcler® applying a standard SYBR green protocol. GAPDH was quantified as a reference-gene for normalization of other genes using the ΔCp-method.

### Immunoblotting

Protein levels were evaluated by immunoblot analysis. Varying numbers of cells were lysed (250,000 MDM, 500,000 HFF/ARPE-19/HeLa and 1,000,000 293T) using lysis buffer supplemented with 1x protease inhibitor and 1% Triton X-100 for 20 min at 4°C. Samples were then stored at −20°C. For SDS-PAGE (polyacrylamide gel electrophoresis) lysates were thawed and kept on ice. Lysates were centrifuged for 10 min at 10,000 rpm at 4°C. Supernatant was mixed 1x SDS-cracking buffer and heated for 10 min at 95°C. Samples were loaded into a 12 % SDS-gel and separated at 80 Volts. Proteins were transferred to a methanol-activated PVDF-membrane (Polyvinylidene fluoride) by wet-blotting for 90 min at 80 Volts. Membranes were then blocked using 5 % milk in TBS for 1 h at RT. Primary antibodies (mouse SAMHD1, 1:500; rabbit pSAMHD1, 1:1000; mouse-anti-Actin, 1:1000) were diluted in 5 % milk in TBS-T (with 0.1% Tween 20) and membranes were incubated overnight at 4°C or for 2 h at RT. Excess antibodies were removed by washing three times with TBS-T and then the membranes were incubated in secondary antibody-solution (IRDye 680 Goat-antiMouse, 1:15,000 or IRDye 800 Goat-anti-Rabbit, 1:15,000 in TBS-T) for 1 h at RT. After washing three times with TBS-T membranes were recorded using FcOdysee. Data was captured and band-intensities were quantified using ImageStudio™lite.

### Software and statistics

For experimental set-up and data analysis Microsoft Word and Excel (Microsoft) were used. Graph-Pad Prism 9 was used for graph generation and statistical analysis. All figures were generated with CorelDrawX7. Other software used included Gen5 v.2.09 (Biotek), Fiji (ImageJ), for image acquisition and analyses, MACS Quantify (Miltenyi) and Flowlogic (Inivai) for flow cytometry, ImageStudio™lite (LiCor) for immunoblotting, LightCycler v.4.1 (Roche) for RT-qPCR, ICE software for the MTT assay.

## RESULTS

### Kinetic of HCMV-mediated SAMHD1 reduction and protein phosphorylation

HCMV uses different mechanisms to counteract SAMHD1. In detail, a transcriptional block of SAMHD1 mRNA synthesis, SAMHD1 degradation and induction of SAMHD1-phosphorylation were observed^29^. Since a complete viral replication cycle in MDMs takes 6 days to be completed, our goal was to explore the kinetics of the different viral counteraction mechanisms. To investigate this, primary human monocyte-derived macrophages (MDMs) were infected with HCMV TB40/E. Cells were harvested 1, 2, 4 or 6 days post infection(dpi) and SAMHD1 and pSAMHD1 levels were measured using flow cytometry and the gating strategy was based on the VPX+ VLP control^29^ (Fig. 1). pSAMHD1 levels were analyzed in three different populations; mock, which are cells treated only with medium, bystanders, which are cells that have been challenged with virus but did not get productively infected and the productively infected cells, which is reflected by immediate early 1 and 2 (IE1/2) expression. SAMHD1 steady-state levels in the infected population of MDM were only slightly reduced 1 dpi as compared to mock and bystander cells and the extent of SAMHD1 reduction increased up to 6 dpi (Fig. 1A). Of note, phosphorylation of SAMHD1 at T592 was already significantly induced as early as one day post infection (average % of pSAMHD1 positive cells; mock: 5.43%, bystanders: 3.35%, infected: 20.34%) with this effect remaining robust throughout all measured time points (Fig. 1B). Hence, HCMV-induces SAMHD1-phosphorylation immediately upon infection whereas SAMHD1 reduction and block of de novo synthesis becomes fully effective at later stages of infection.

**Fig. 1:**
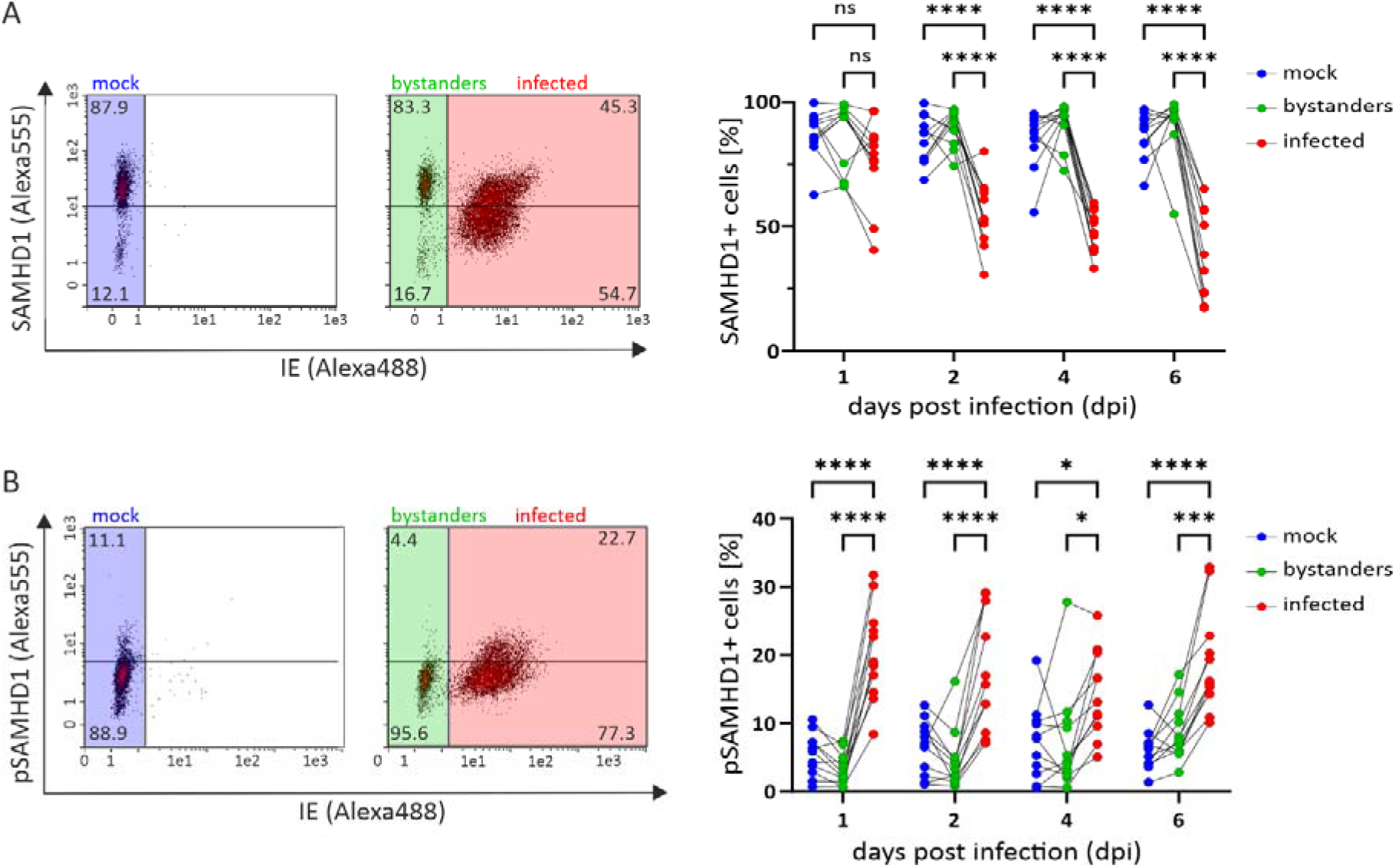
(p)SAMHD1 kinetics upon HCMV infection. MDMs were infected with HCMV TB40E and harvested after 1,2,4 or 6 days. Cells were intracellularly stained for SAMHD1 (A) or pSAMHD1 (B) and IE to discriminate infected cells and analyzed with flow cytometry. Primary flow cytometry data of one representative donor are depicted and indicate the gating strategy. Each three connected dots represent one donor (n=11). Significance was tested with two-way analysis of variance (ANOVA) with Tukey’s correction. ****, P<0.0001; ***, P<0.001; **, P<0.01; *, P<0.05.

### Induction of SAMHD1 phosphorylation is a conserved feature of various primary HCMV clinical isolates

The ability of HCMV to phosphorylate SAMHD1 was shown for HCMV-lab adapted strains only and it is currently unclear, if this is a conserved feature of primary, clinical HCMV strains. In order to assess if HCMV clinical isolates retain the ability to phosphorylate SAMHD1, we tested a number of fibroblast-tropic HCMV strains on human foreskin fibroblasts (HFFs). As compared to macrophages, HFF have much lower SAMHD1 levels and induction of pSAMHD1 is less pronounced^49^. HFF were infected with HCMV TB40/E WT, TB40/E-delUL16-GFP or 5 different clinical isolates (Fig. 2). After 48 hours, cells were harvested and SAMHD1 and pSAMHD1 levels were detected by flow cytometry. In HFF, in agreement to previous observations^49^, the low SAMHD1 steady-state levels increased upon HCMV infection which was consistent within the lab strain TB40/E and the 5 clinical isolates used (Fig. 2A). The increase in the infected population was between 12- to 17-fold compared to mock, with a similar increase in SAMHD1 levels when comparing the bystanders to mock cells (5- to 16-fold). More importantly, the 5 different clinical strains induced pSAMHD1 (3.5- to 11-fold more pSAMHD1-positive cells in infected than in mock populations) to a similar extent as the lab-adapted TB40 strains (TB40/E-WT: 4.4-fold, TB40E-GFP: 8.0-fold increase of pSAMHD1-positive cells) (Fig. 2B). In addition, we used an endothelial/macrophage-tropic clinical isolate to infect MDMs and measured the SAMHD1 and pSAMHD1 levels after 1, 2, 4 or 6 days of infection. As a reference, we used the TB40/E WT strain. The clinical isolate as well as TB40/E exhibited a similar tendency in the kinetics of SAMHD1 reduction (Fig. 2C) and pSAMHD1 induction (Fig. 2D). Together, these data indicate that the ability to phosphorylate, and hence inactivate the antiviral host factor SAMHD1, is a conserved function of different primary clinical HCMV isolates.

**Fig. 2:**
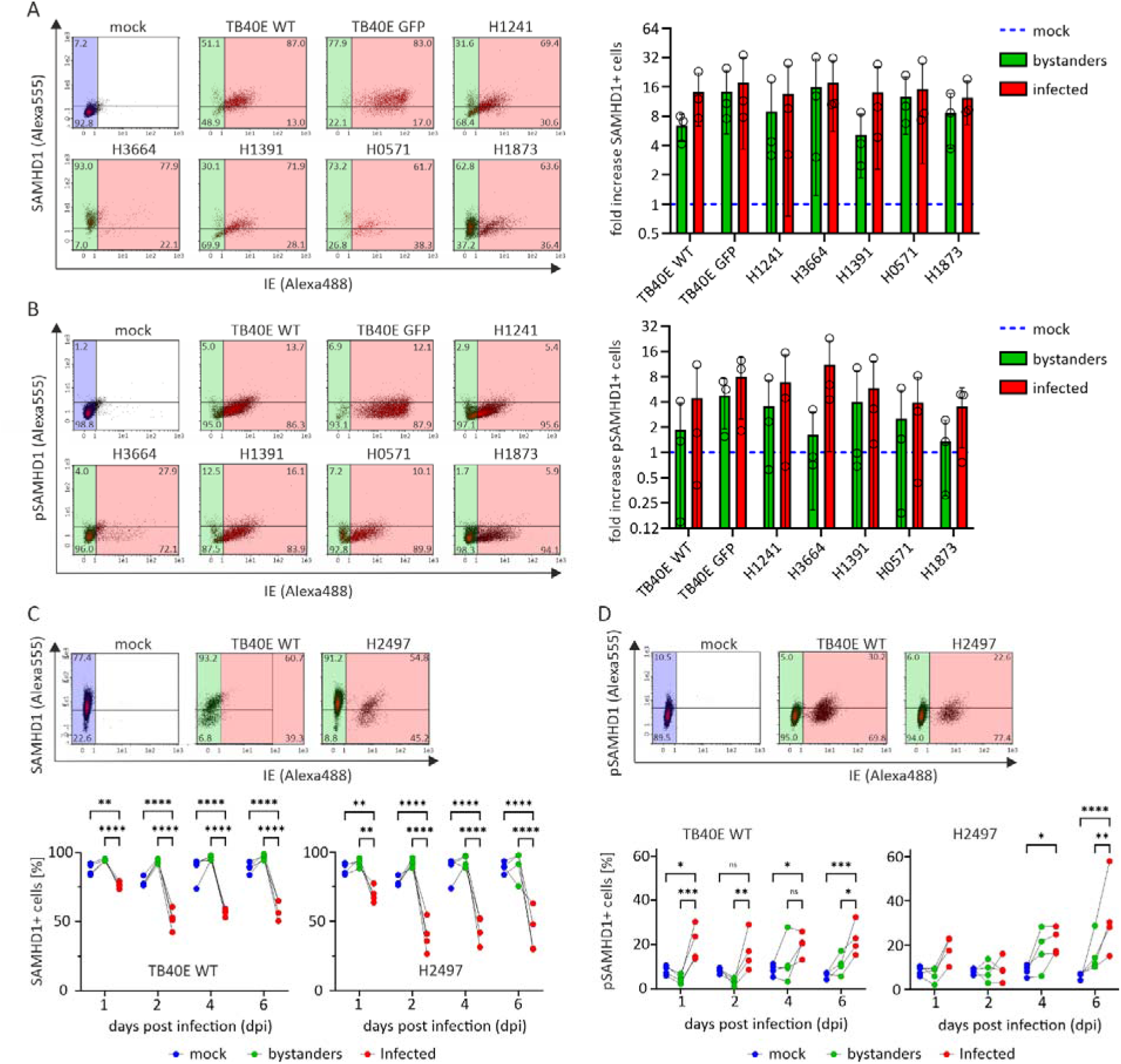
Clinical HCMV isolates retain the ability to phosphorylate SAMHD1. HFFs were infected with a panel of clinical isolates and TB40/E WT and TB40/E-delUL16-GFP as a reference. Cells were harvested 2 dpi and intracellularly stained for SAMHD1 (A) and pSAMDH1 (B), prior to flow cytometry analysis. MDMs were infected with a macrophage tropic clinical isolate or with TB40E WT as a reference and harvested 1,2,4 or 6 dpi. MDMs were then intra-stained for SAMHD1 (C) or pSAMHD1 (D) and measured using flow cytometry. Primary flow cytometry data of one representative donor for the macrophages (n=4) (C, D left), or one representative replicate for the HFFs (n=3) (A, B left) are shown. Each three connected dots represent one donor. Significance was tested with two-way analysis of variance (ANOVA) with Tukey’s correction. ****, P<0.0001; ***, P<0.001; **, P<0.01; *, P<0.05; ns: not significant.

### CDK 4/6 inhibitors suppress HCMV replication in MDMs

HCMV infection activates the host kinase CDK2 by increasing the levels of its interaction partner cyclin E^33^. The cyclin E/CDK2 complex phosphorylates, among others, SAMHD1^20^. Furthermore, the HCMV encoded kinase and CDK-homologue pUL97 phosphorylates SAMDH1 to counteract the antiviral activity of SAMHD1^29^. We hence hypothesized that the FDA-approved CDK4/6 inhibitors Palbociclib (PC), Ribociclib (RC) and Abemaciclib (AC) might inhibit HCMV-replication in macrophages. As a reference, we used the current standard of care against HCMV, Ganciclovir (GCV), which inhibits viral genome replication. MDMs were pre-treated with the different compounds for 16 hours and infected with TB40/E WT. After 6 days of infection, supernatants of mock or HCMV infected MDMs were collected and back-titrated on HFFs, to determine the overall potency of virus replication in the presence of the antiviral drugs (Fig. 3). All three CDK4/6 inhibitors exhibited antiviral activity with AC being most potent with an IC50 of 42 nM, which was even superior to the positive control Ganciclovir (IC50: 118 nM) (Fig. 3A and B, table 1). In addition, AC showed no cytotoxicity in the working concentrations with a CC50 of 14.7 μM (Fig. 3B), resulting in a therapeutic index (CC50/IC50) of 350 (Table 1). Of note, AC and the other CDK4/6 inhibitors were not active against HCMV replication in HFFs (Supplemental Fig. 1), presumably due to the fact that that HFFs express low levels of SAMHD1 (Supplemental Fig. 2), agreeing with previous findings that Palbociclib was antivirally active only in SAMHD1 expressing cells^44^. In sum, FDA approved CDK4/6 inhibitors are highly potent in suppressing HCMV replication in macrophages with AC showing superior antiviral activity as compared to GCV or other CDKIs.

**Fig. 3:**
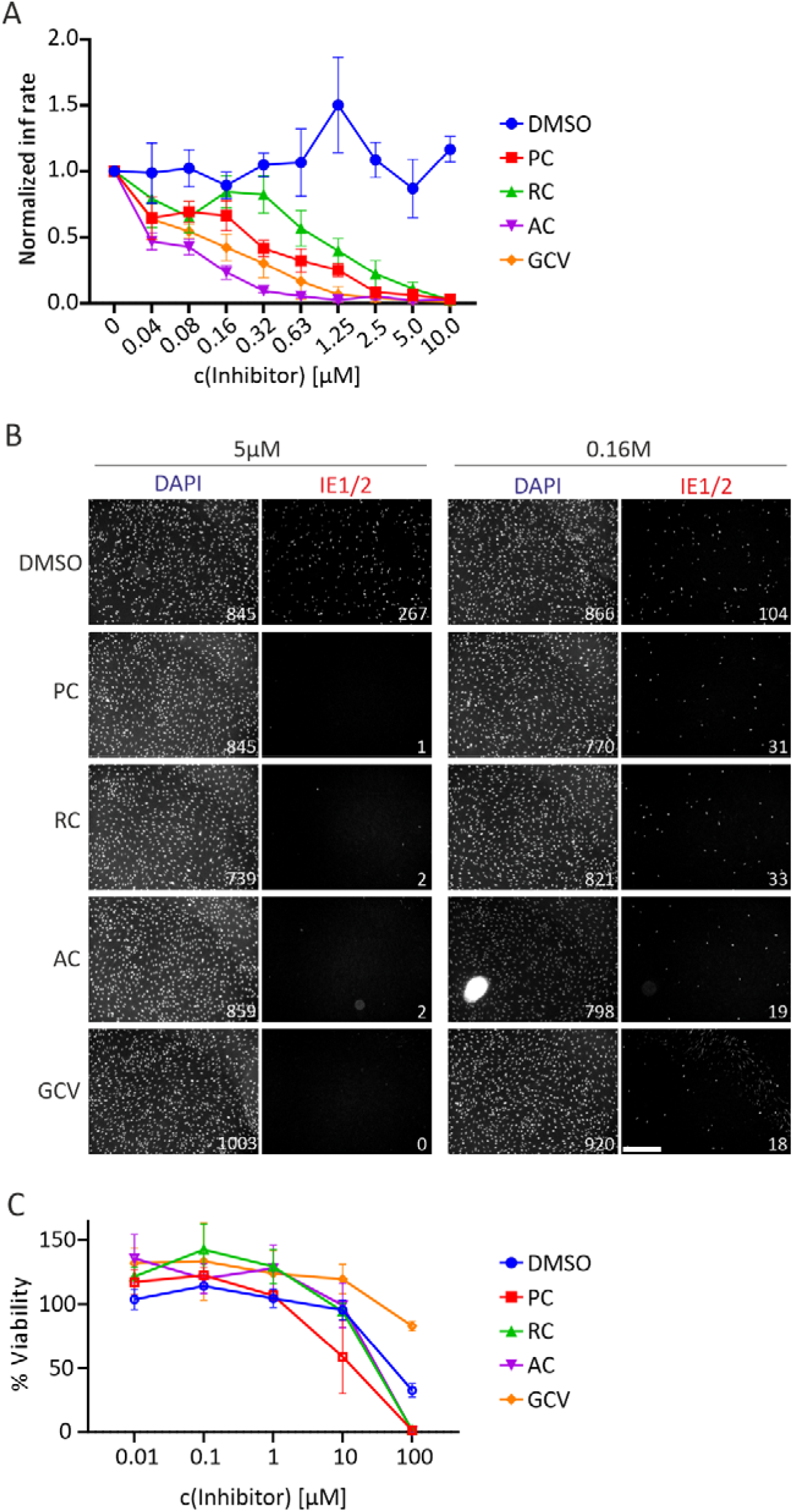
Abemaciclib robustly blocks HCMV replication with low cytotoxicity in primary human macrophages. MDMs were pre-incubated with serial dilutions of a panel of CDK4/6 inhibitors for 16 hours. Cells were then infected with HCMV TB40/E and treated again with CDK inhibitors or DMSO control for 6 days. At 6 dpi supernatants were collected and used to infect HFFs for 16 hours. Cells were fixed and intracellularly stained for the viral proteins IE 1/2. Subsequently, nuclei were stained with DAPI and infection rates were analyzed by immunofluorescence. The IC50s of each compound (A) were calculated using GraphPad Prism 9. Representative images of HFFs treated with different inhibitors and infected with HCMV. Scale bar, 0.5 mm (B). MTT assay was performed to determine the cytotoxicity of each inhibitor (C). Error bars show SEM, n=3. PC: Palbociclib; RC: Ribociclib; AC: Abemaciclib; GCV: Ganciclovir

**Table 1.** IC50, CC50 and therapeutic indices (TI) of a panel of different CDK inhibitors against HCMV replication in primary macrophages (n=3 or 4).

| Inhibitor | Target | IC50 (nM) | CC50 (μM) | Therap. Index |
| --- | --- | --- | --- | --- |
| Palbociclib | CDK4, CDK6 | 355 | 12.3 | 34.6 |
| Ribociclib | CDK4, CDK6 | 1294 | 20.2 | 15.6 |
| Abemaciclib | CDK4, CDK6 | 42 | 14.7 | 350 |
| K03861 | CDK2 | 1028 | 1.28 | 1.25 |
| LDC4297 | CDK7 | 37 | 1.22 | 33 |
| JSH-150 | CDK9 | 223 | 0.31 | 1.4 |
| Ganciclovir | - | 118 | >100 μM | N/A |

### Abemaciclib blocks HCMV-mediated phosphorylation of SAMHD1 by targeting pUL97

The virally encoded kinase pUL97 is a CDK-homologue and phosphorylates SAMHD1. We hence hypothesized that the CDKIs might inhibit HCMV-mediated SAMHD1 phosphorylation, which could be a potential mode of antiviral action. For this, HCMV-infected MDMs were treated with different CDKIs and levels of total and phosphorylated SAMHD1 were measured at different time points post infection (Fig. 4). Palbociclib, in agreement with a previous report^45^, blocked the phosphorylation of SAMHD1 in infected cells at 2 dpi down to 8.7% compared to 17.7% in the DMSO control, while Ribociclib, the inhibitor with the lowest antiviral activity, didn’t result in reduction of pSAMHD1 levels (18.6% pSAMHD1-positive cells) similar to the negative controls that were just DMSO treated or treated with GCV (Fig. 4A,B). Abemaciclib blocked the HCMV-mediated phosphorylation of SAMHD1 significantly from day 2 on (4.5% pSAMHD1-positive cells) with nearly no detectable levels of pSAMHD1 at 4 dpi (Fig. 4B).

**Fig. 4:**
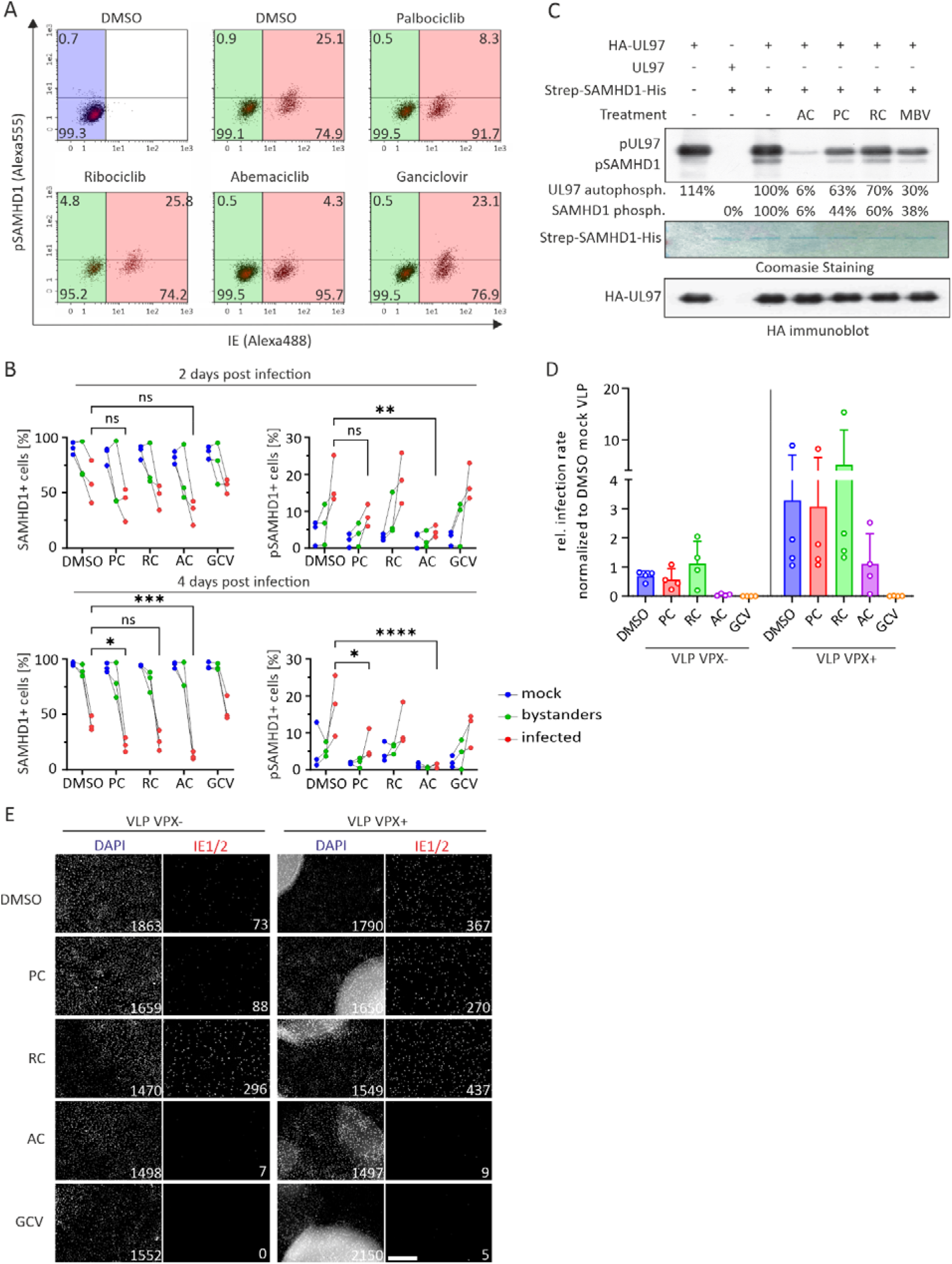
Abemaciclib blocks the phosphorylation of SAMHD1 by inhibiting the activity of the viral kinase pUL97. MDMs were pre-treated with CDK4/6 inhibitors, or DMSO and infected with HCMV TB40/E. After virus removal, cells were re-treated with CDK4/6 inhibitors or DMSO or Ganciclovir and harvested and fixed at 2 or 4 dpi. Subsequently, they were stained for pSAMHD1 and analyzed with flow cytometry (A, B). Each three connected dots represent one donor (n=3). Significance was tested with two-way analysis of variance (ANOVA) with Tukey’s correction. ****, P<0.0001; ***, P<0.001; **, P<0.01; *, P<0.05. The indicated kinase inhibitors were added to in vitro kinase assays measuring the activity of HA-immunopurified pUL97 by pUL97 auto-phosphorylation and SAMHD1 phosphorylation (C). A non-tagged version of pUL97 served as control. Input levels were controlled by Coomassie staining (SAMHD1) and HA-immunoblotting (UL97). MDMs were pretreated with no VLP, VLP VPX- or VLP VPX+ in order to knock down SAMHD1. Two hours later, cells were infected, and supernatants were collected 6 days post infection. The supernatants, in turn, were used to infect HFFs for 16 hours. Cells were fixed and stained for DAPI and IE1/2 before being analyzed by immunofluorescence. Error bars show SEM (D). Representative pictures of cells treated with either VPX+ or VPX-VLPs and infected with TB40E. Scale bar, 0.5 mm (E). PC: Palbociclib; RC: Ribociclib; AC: Abemaciclib; GCV: Ganciclovir. Significance was tested with two-way analysis of variance (ANOVA) with Tukey’s correction. ****, P<0.0001; ***, P<0.001; **, P<0.01; *, P<0.05; ns: not significant.

Formally, as HCMV hijacks cellular kinases, lower levels of pSAMHD1 could be due to inhibition of cellular CDKs or the virally encoded kinase pUL97, that is known to phosphorylate SAMHD1^29^. We therefore tested in an *in vitro* kinase assay the potency of the different CDKIs to inhibit pUL97 autophosphorylation and the pUL97-mediated phosphorylation of SAMHD1 (Fig. 4C). Of note, Abemaciclib was a highly potent inhibitor of pUL97 autophosphorylation as well as SAMHD1 phosphorylation (down to 6% compared to 100% in DMSO) which was in stark contrast to the direct inhibitory activity of the other CDKIs and Maribavir, which is used as a specific pUL97-inhibitor. These findings suggest that Abemaciclib’s antiviral activity is due to potent inhibition of pUL97 and its ability to phosphorylate SAMHD1. Of note, when SAMHD1 was knocked down using the accessory SIV protein VPX, Abemaciclib retained some of its antiviral activity (Fig. 4D and E). This indicates that SAMHD1-independent mechanisms play an additional role in Abemaciclib’s antiviral activity.

### The CDK7 inhibitor LDC4297 is a potent inhibitor of HCMV replication in MDMs and robustly suppresses SAMHD1 phosphorylation

Abemaciclib, the most active of the 3 different CDK4/6 inhibitors tested, has inhibitory activity against other CDKs, as in particular CDK7^50,51^ which is the closest cellular homologue of pUL97. We hence hypothesized that specifically CDK7-targeting CDKis could be potent inhibitors of HCMV replication and pUL97-mediated SAMHD1 phosphorylation. For this we tested inhibitors of CDK2 (K03861), CDK7 (LDC4297) and CDK9 (JSH-150) for antiviral activity in MDM (Fig. 5). LDC4297, a CDK7 inhibitor which is already known for its anti-HCMV effect in HFFs^41^, showed potent inhibition of HCMV replication in primary macrophages (IC50: 37 nM, CC50: 1.22 μM, therapeutic index: 33) (Fig. 5A and B, table 1). This activity was much more pronounced as compared to the CDK2 inhibitor K03861 and superior to the antiviral activity of JSH-150, which was furthermore toxic to MDM at relatively low concentrations (CC50: 0.31 μM) (Fig. 5C, table 1). Of note, in line with our data, LDC4297 is highly superior in inhibiting phosphorylation of SAMHD1 when compared to K03861 (Fig. 5D). In conclusion, inhibition of SAMHD1-phosphorylation by specific CDKIs that are FDA-approved or in advanced clinical development, is a potent strategy to inhibit HCMV-replication in primary human macrophages.

**Fig. 5:**
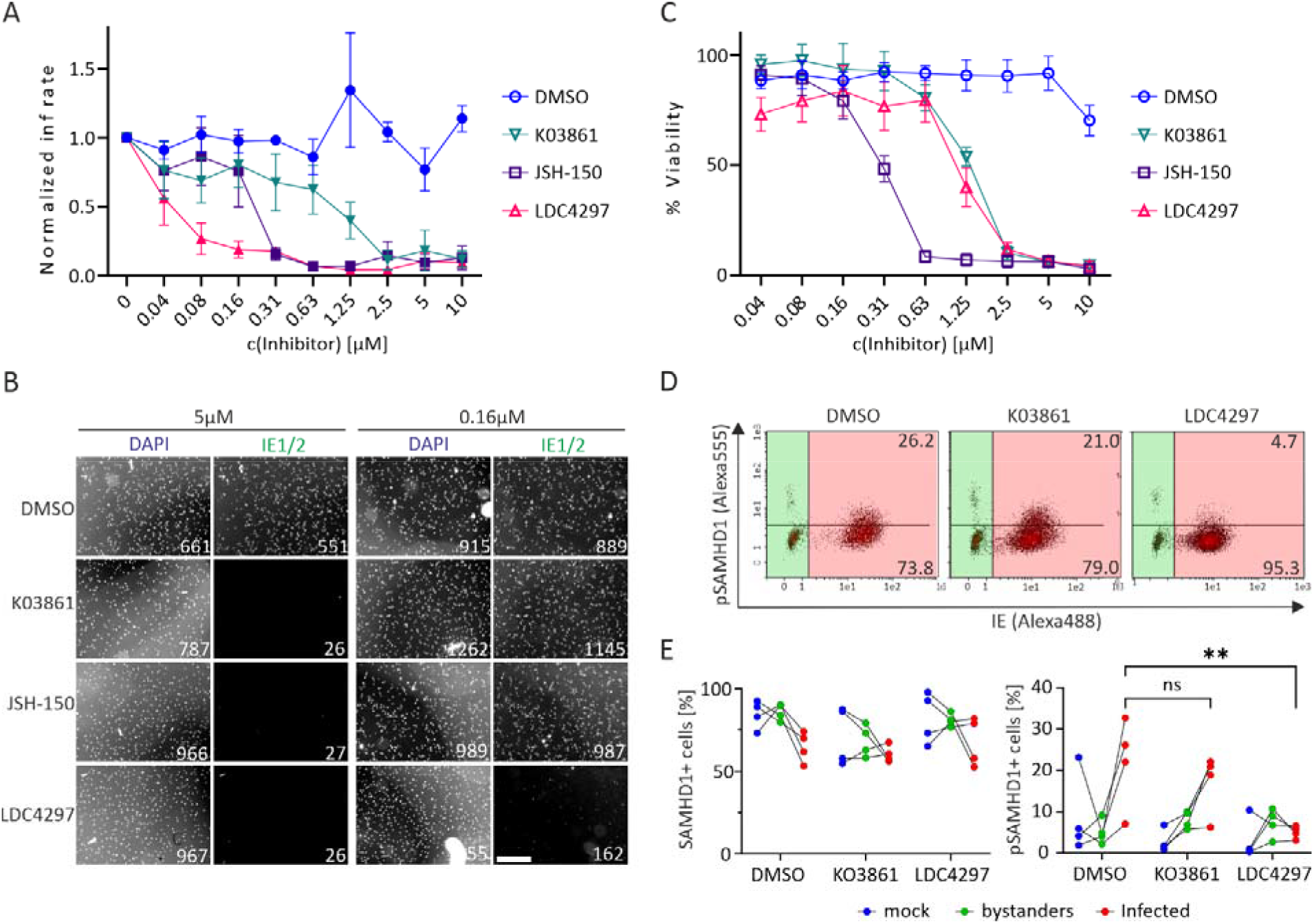
A CDK7 inhibitor, LDC4297 exhibits suppression of HCMV replication in primary human macrophages. MDMs were infected with HCMV TB40/E and then treated with serial dilutions of CDK inhibitors (K03861: CDK2 inhibitor, JSH-150: CDK9 inhibitor, LDC4297: CDK7 inhibitor) or DMSO control for 6 days. At 6 dpi, supernatants were collected and used to infect HFFs for 16 hours. Cells were fixed and intracellularly stained for the viral proteins IE1/2. Subsequently, nuclei were stained with DAPI and infection rates were analyzed by immunofluorescence microscopy (A). Representative images of HFFs treated with different inhibitors and infected with HCMV. Scale bar, 0.5 mm (B). MTT assay was performed to determine the cytotoxicity of each inhibitor (C). MDMs were infected with HCMV TB40/E, then treated with CDK inhibitors and harvested and fixed at 2 dpi. Subsequently, they were stained for pSAMHD1 and analyzed with flow cytometry (D, E). Primary flow cytometry data of one representative donor are depicted, indicating the gating strategy (D). Each three connected dots represent one donor (n=4) (E). Error bars show SEM. The significance was tested by a two-way analysis of variance (ANOVA) with Tukey’s correction. **, P<0.01; ns: not significant.

## DISCUSSION

HCMV uses kinases to optimize its viral genome replication and evade the antiviral immune response. Amongst others, HCMV induces phosphorylation of the host cell restriction factor SAMHD1 through the virus encoded kinase pUL97, resulting in SAMHD1 inactivation^29^. Moreover, HCMV antagonizes SAMHD1 by at least two additional mechanisms, that are proteasomal degradation and transcriptional suppression of SAMHD1^29^. The relative importance of these SAMHD1 inactivating mechanisms for HCMV replication were previously unclear. Analyzing the temporal order of pSAMHD1 and total SAMHD1 we found that phosphorylation takes place rapidly and is already evident as soon as 24 hours post infection. In contrast total SAMHD1 levels remained unchanged, with a reduction in SAMHD1 levels that was only evident later during infection.

This finding indicates that early phosphorylation of SAMHD1 by pUL97 is essential for HCMV to establish productive replication in macrophages. The relevance of HCMV-mediated SAMHD1 phosphorylation is further corroborated by the highly conserved ability of primary clinical HCMV isolates to induce pSAMHD1 in HFF and macrophages. We hence hypothesized that suppression of SAMHD1-phosphorylation could prove as a highly effective antiviral approach.

During the cell cycle, phosphorylation of SAMHD1 is controlled by CDKs, especially CDK1 and CDK2^19^. CDK2 in turn is active only when it is phosphorylated, by CDK6^17^. HCMV is known to hijack CDKs and selectively activate CDK/cyclin complexes^33^. Palbociclib (PC), a CDK4/6 inhibitor blocks the phosphorylation of SAMHD1 in primary macrophages in a CDK6/CDK2 manner, in the context of HIV-1 infection^45^. Furthermore, pUL97 is a CDK-homologue^52,53^. This prompted us to evaluate the antiviral activity of 3 FDA approved CDK4/6 inhibitors. The CDKis had a highly divergent pattern of antiviral activity, with PC and Ribociclib (RC) inhibiting HCMV replication in macrophages with modest to low efficacy, whereas Abemaciclib (AC) showed superior antiviral activity, which even outcompeted the gold standard in HCMV treatment, Ganciclovir. Anti-HCMV activity of AC has been investigated recently in HFF, showing limited antiviral activity with an IC50 of ~8 μM^39^, which is approximately 200-fold higher as reported by us in macrophages. In line, AC showed no to low anti-HCMV activity in HFF that express no to low levels of SAMHD1. This suggests that SAMHD1 is the major target of antivirally active CDK4/6 inhibitors and is in agreement with a study showing that PC only exerts activity against Herpes Simplex Virus Type I in SAMHD1 expressing cells^44^. In line, only AC, but not PC or RC was a highly potent inhibitor of SAMHD1-phosphorylation in HCMV infected cells. A finding corroborating SAMHD1 to be the main responsible therapeutic target of AC. To further analyze this, we performed in vitro kinase assays (IVKAs) and clearly demonstrated that AC is a highly potent pUL97 inhibitor, directly suppressing pUL97 autophosphorylation and pUL97-mediated SAMDH1-phosphorylation. This could be due to ACs ability to block the ATP-cleft of various CDKs^50^. AC’s anti-pUL97 activity was even higher compared to Maribavir, which was developed as specific pUL97 inhibitor^54^. Inhibition of pUL97 and its various downstream targets furthermore explains the SAMHD1-independent antiviral effects of AC, that are measurable upon SAMHD1-depletion from infected macrophages.

Although AC robustly blocks pUL97 activity, it is a CDK4 and CDK6 inhibitor, while pUL97 has functional overlaps with CDK-family members CDK1, CDK7 and CDK9^55^. We therefore hypothesized that other CDK inhibitors which more specifically block the CDK family orthologs to pUL97 could exert high antiviral activity. However surprisingly, the CDK9 inhibitor JSH-150 was toxic in macrophages and in lower concentrations did not show pronounced antiviral activity, similar to the CDK2 inhibitor K03861. On the other hand, the CDK7 inhibitor LDC4297 had an IC50 comparable to AC, but a lower therapeutic index as it exhibited cytotoxicity already at lower concentrations. Again, similar to AC, LDC4297 suppresses the phosphorylation of SAMHD1, but not the CDK2 inhibitor K03681. It is exciting and noteworthy that LDC4297 and AC have been suggested to exert anti-HCMV activity in a highly synergistic manner^39^. This is explainable by our data establishing AC as a novel and highly potent inhibitor of pUL97 and the fact that LDC4297 doesn’t affect the kinase activity of pUL97^41^. Hence, both inhibitors suppress SAMHD1-phosphorylation and thereby HCMV replication by different and apparently complementary modes-of-action.

In summary, in this study we demonstrate how AC, a CDK4/6 inhibitor, already applied as a treatment against breast cancer^56^ can be used as an antiviral drug against HCMV. AC is active in low concentrations with an IC50 in the nanomolar range in macrophages. It targets the viral kinase pUL97 and thereby downstream phosphorylation of its effectors including the antiviral factor SAMHD1, which emerges as highly relevant druggable target. AC could be combined with drugs which use different modes of action (like it has been already shown for combination of AC with LDC4297^39^), to achieve synergistic effects, meaning maximum antiviral potential and minimum side-effects. Such approaches could be promising alternatives to the current therapeutic strategies against HCMV.

## Consent for publication

All authors gave their consent to publish, All authors read and approved the final manuscript.

## Availability of data and material

All data generated and analyzed during this study are included in this published manuscript.

## Authors’ contributions

GVS, MF, AD and RB performed most of the experiments, in detail HCMV infection of MDM, antiviral activity of CDKIs, FACS-analysis for SAMHD1 and pSAMHD1 levels and qRT-PCR. IG and LW panned and conducted the IVKA. KH provided patient isolated HCMV strains. GVS and MS planned and designed most of the experiments and analyzed the data. GVS and MS wrote the initial manuscript draft. MS conceived and and supervised the overall study and provided resources. All authors contributed to editing and developed the manuscript to its final form.

## Acknowledgements

We thank the team of the Transfusion Medicine of the University Hospital Tübingen for their support in providing buffy coat to differentiate primary macrophages and all anonymous blood donors for their support.

## Supplemental figures

**Fig. S1:**
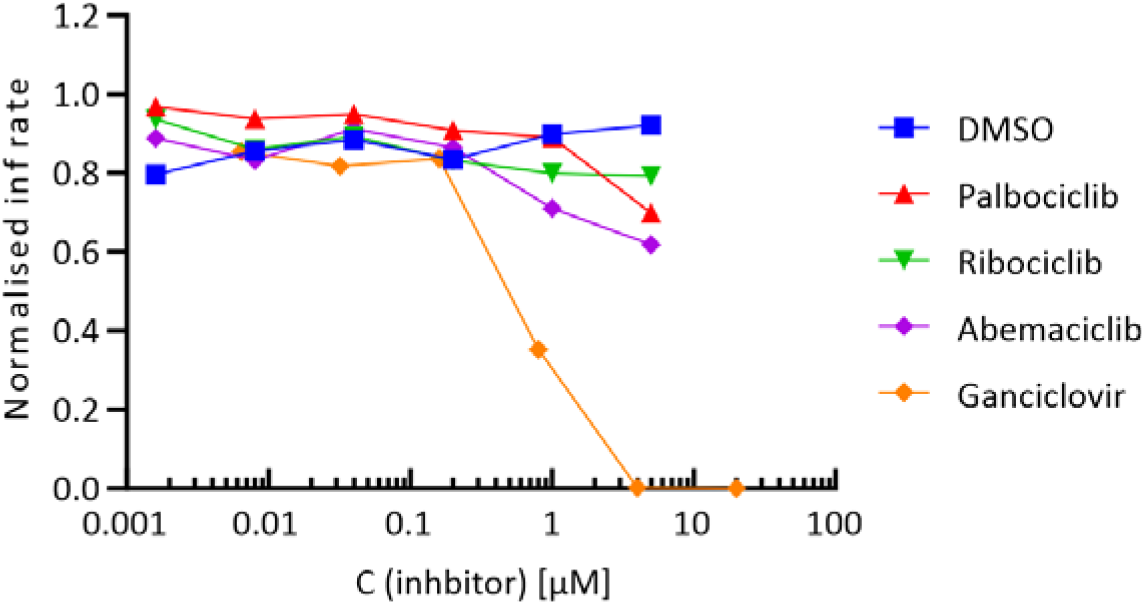
CDK4/6 inhibitors do not block HCMV replication in human foreskin fibroblasts. HFFs (n=1) were pre-incubated with serial dilutions of a panel of CDK4/6 inhibitors for 16 hours. Cells were then infected with HCMV TB40E and treated again with CDK inhibitors or DMSO control for 6 days. At 6 dpi supernatants were collected and used to infect fresh HFFs for 24 hours. Cells were fixed and intra-stained for the viral proteins IE 1/2. Subsequently, nuclei were stained with DAPI and infection rates were analyzed by immunofluorescence.

**Fig. S2:**
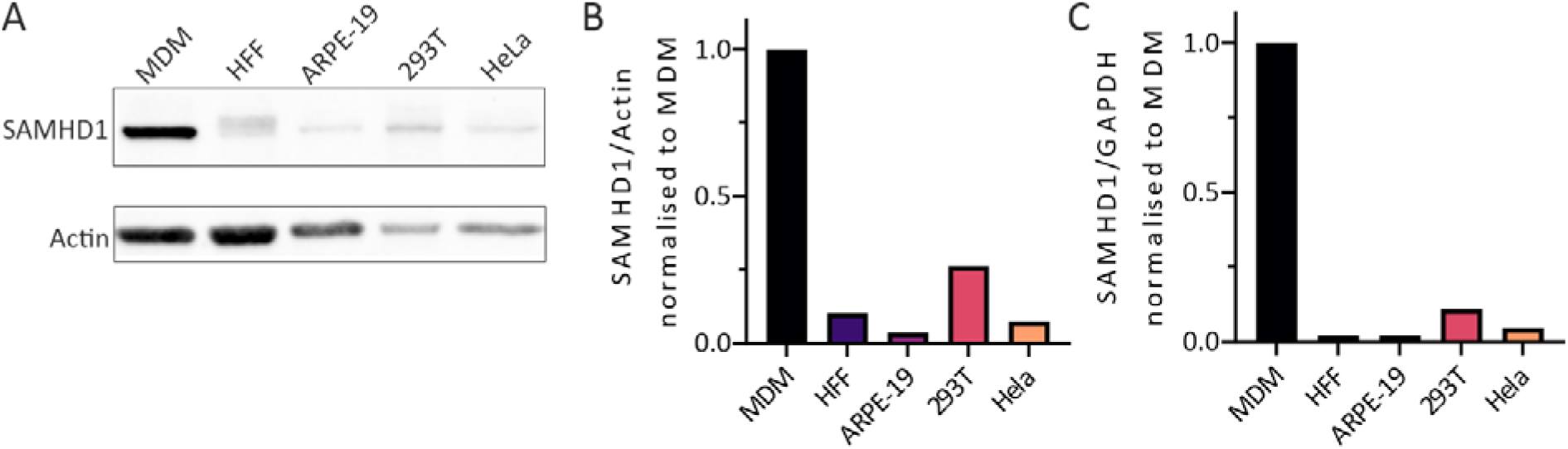
SAMHD1 steady-state levels in different cell types. A panel of various cell types (MDM, HFF, ARPE-19, 293T HEK, HeLa) (A, B) were lysed and analyzed by immunoblotting for SAMHD1 (MW: 72 kDa) and actin (MW: 42 kDa) as a loading control. (C) RNA was extracted from the same cells and reversed transcribed to cDNA, before amplifying SAMHD1 and GAPDH as a housekeeping gene control, with the appropriate primers. Signals were detected using the RT-qPCR SYBR-Green method.

